# Recombination map tailored to Native Hawaiians improves robustness of genomic scans for positive selection

**DOI:** 10.1101/2023.07.12.548735

**Authors:** Bryan L. Dinh, Echo Tang, Kekoa Taparra, Nathan Nakatsuka, Fei Chen, Charleston W. K. Chiang

## Abstract

Recombination events establish the patterns of haplotypic structure in a population and estimates of recombination rates are used in several downstream population and statistical genetic analyses. Using suboptimal maps from distantly related populations may reduce the efficacy of genomic analyses, particularly for underrepresented populations such as the Native Hawaiians. To overcome this challenge, we constructed recombination maps using genome-wide array data from two study samples of Native Hawaiians: one reflecting the current admixed state of Native Hawaiians (NH map), and one based on individuals of enriched Polynesian ancestries (PNS map) with the potential to be used for less admixed Polynesian populations such as the Samoans. We found the recombination landscape to be less correlated with those from other continental populations (e.g. Spearman’s rho = 0.79 between PNS and CEU (Utah residents with Northern and Western European ancestry) compared to 0.92 between YRI (Yoruba in Ibadan, Nigeria) and CEU at 50 kb resolution), likely driven by the unique demographic history of the Native Hawaiians. PNS also shared the fewest recombination hotspots with other populations (e.g. 8% of hotspots shared between PNS and CEU compared to 27% of hotspots shared between YRI and CEU). We found that downstream analyses in the Native Hawaiian population, such as local ancestry inference, imputation, and IBD segment and relatedness detections, would achieve similar efficacy when using the NH map compared to an omnibus map. However, for genome scans of adaptive loci using integrated haplotype scores, we found several loci with apparent genome-wide significant signals (|Z-score| > 4) in Native Hawaiians that would not have been significant when analyzed using NH-specific maps. Population-specific recombination maps may therefore improve the robustness of haplotype-based statistics and help us better characterize the evolutionary history that may underlie Native Hawaiian-specific health conditions that persist today.

## Introduction

Knowledge of the recombination landscape across the genome informs patterns of haplotypic structure in a population and is used in several population and statistical genetic analyses such as imputation, local ancestry inference (LAI), identity-by-descent (IBD) inference, and genomic scan of adaptive loci, among others. Fine-scale differences in recombination landscapes are known to exist between populations (Hassan et al., 2021; Hinch et al., 2011; Spence & Song, 2019a; van Eeden et al., 2022; Wegmann et al., 2011). For example, recombination maps for HapMap populations CEU (Utah residents of predominantly Northern and Western European ancestries) and YRI (Yoruba in Ibadan, Nigeria) correlate well at large resolutions but have poorer correlation at finer scales (Wegmann et al., 2011). The deCODE map, based on pedigrees of individuals with European-ancestries from Iceland, correlates better with CEU than YRI (Wegmann et al., 2011). Any differences in recombination landscapes between populations can be due to both the divergent demographic histories as well as genetic differences in usage of recombination hotspots (Hinch et al., 2011). However, we have limited understanding of the recombination landscape for populations not represented by the 1000 Genomes Project (1KGP) (Auton et al., 2015), although efforts in developing these maps are starting for diverse populations such as the Nama people of southern Africa (van Eeden et al., 2022). Because large-scale pedigree data are rarely available for diverse populations, most attempts to characterize the recombination landscape of a population begin with population-scaled rates or their derivatives.

Given the prevalent use of recombination maps in genetic analysis and the fine-scale differences between populations, it may be suboptimal to use a recombination map from a distantly related population. This underdevelopment of available genomic resources, in general, may further exacerbate the known disparity in transferring genomic insights and downstream benefits to populations relatively distant from the study populations. Furthermore, the utility of a population specific recombination map on downstream analyses has not been extensively explored and studies have reached different qualitative conclusions. For an isolated, homogenous population like the Finns (Hassan et al., 2021), a population-specific map did not show a large impact on downstream analyses such as phasing and imputation, although this could be due to general haplotypic similarity between Finns and the abundance of European-ancestry samples represented in 1000 Genomes or that information from recombination plays only a secondary role in the specific analyses evaluated. On the other hand, population-specific recombination maps have also been shown to impact haplotype-based scans of positive selection in the Nama (van Eeden et al., 2022), a population not adequately represented by available maps which emphasize West African ancestry.

In this work, we focused on characterizing the recombination landscape for the Native Hawaiians, using genome-wide genotyped individuals from the Multiethnic Cohort (MEC) Study (Kolonel et al., 2000). Native Hawaiians are an admixed, indigenous, and underrepresented population that, with a current population of 680,353 individuals, account for only 0.2% of the U.S. population (*U.S. Census Bureau Releases Key Stats in Honor of 2023 Asian American, Native Hawaiian, and Pacific Islander Heritage Month*, 2023). The lack of resources to advance genomic research among Native Hawaiians has previously been detailed (Chiang, 2021; Lin et al., 2020). We thus also explored the impact of a population-specific map on multiple downstream analyses for the Native Hawaiians. The lack of available whole genome sequencing data and the relatively small cohort size limited the applicable methods that could be used to accurately infer a recombination for this population. We used LDhat (Auton & McVean, 2007) to estimate the recombination landscape. LDhat models the observed linkage disequilibrium (LD) between pairs of SNPs through a Bayesian reversible-jump Markov chain Monte Carlo process to estimate the population recombination rate across the genome and has been shown to infer accurate recombination maps using array data with good concordance to maps inferred through other methods (Spence & Song, 2019b; Zhou et al., 2020). Because of European colonization and subsequent waves of immigrations to the Hawaiian islands, we modeled the Native Hawaiians to include African-, East Asian-, and European-related ancestries in addition to Polynesian ancestries following previous work on this population (Lin et al., 2020; Sun et al., 2021). As a result, we created and characterized two recombination maps from our study sample: one based on a random subset of our sample reflecting the current admixed state of Native Hawaiians, and one based on a subset of individuals with enriched Polynesian ancestries. The latter map was constructed to gain insights on the recombination landscape of ancestral Polynesians and potentially provide a viable map for relatively unadmixed Polynesian populations such as the Samoans. We then evaluated the impact of a Native Hawaiian-specific map for downstream analyses such as LAI, imputation, IBD segment inference, and genome-wide scans of adaptation using haplotype-based statistics.

## Materials and Methods

### Study cohort and data

The primary genetic dataset is a subcohort of the Multiethnic Cohort (MEC) Study, which is a prospective epidemiological cohort of >215,000 individuals established as a collaboration between the University of Hawai‘i and University of Southern California (Kolonel et al., 2000). We focused on the 3,940 Native Hawaiian (MEC-NH) individuals genotyped on the Illumina MEGA array as part of the PAGE consortium (Wojcik et al., 2019). We additionally used MEGA array data from 5,325 African Americans (MEC-AA) from the same cohort for specific comparisons in the present study. Detailed descriptions of sample processing and QC can be found in previous publications (Lin et al., 2020; Sun et al., 2021). In short, we restricted our analysis to biallelic SNPs with positions found in 1KGP and genotyped in greater than 95% of the individuals, leaving 1,326,678 starting SNPs in MEC-NH population. We selected two sets of 96 MEC-NH and one set of 96 MEC-AA individuals from these two datasets to construct an LD-based recombination map (below). We then used the remaining individuals for evaluations of the maps in downstream statistical and population genetic applications.

In addition to the primary datasets from the PAGE consortium, we had access to 307 MEC-NH individuals previously genotyped on the Illumina MEGAex array for a study of obesity (Lim et al., 2019) and 453 MEC-NH individuals previously genotyped on the Illumina 660W array for study of breast cancer (Siddiq et al., 2012). For a subset of these NH individuals, we also had targeted exon sequencing data across 746 exons in 37 genes as part of a breast cancer study (Hu et al., 2021); we used these in combination to evaluate the impact of recombination maps on genotype imputation. In total, 607 sequenced individuals overlap with our MEC-NH data: 453 from the NHBC study, 149 from the PAGE study, and 5 from the obesity and PAGE studies.

### Native Hawaiian-specific recombination map using LDhat and IBDrecomb

Recombination maps were inferred using LDhat (Auton & McVean, 2007) with data ascertained on the MEGA array. We modeled the analytic pipeline using LDhat after previously published descriptions on similar efforts to infer the recombination landscape in humans and primates (Spence & Song, 2019a; Xue et al., 2020; Zhou et al., 2020). LDhat relies on lookup tables to allow for tractable computation. Because the creation of lookup tables is computationally taxing, we limited our analysis to the largest table based on 192 haplotypes that was provided with the software. For this reason, we generated two recombination maps: one using 96 individuals most enriched with the Polynesian ancestries that were found predominantly in Native Hawaiians (PNS, proportion of Polynesian ancestries were previously estimated (Lin et al., 2020)), and one using 96 individuals randomly selected from our Native Hawaiian samples reflecting admixtures in our study sample (NH) (Lin et al., 2020; Sun et al., 2021). These subsets were chosen to generate maps representative of a population with predominantly Polynesian ancestries and the current NH population, respectively. We limited our analysis to the autosomes.

We updated the genome build of our data to hg38 using Liftover (Hinrichs et al., 2006) and then phased all available individuals with Eagle (v2.4.1, without including a phased reference) using the default omnibus map based on HapMap populations (International HapMap Consortium et al., 2007; Loh, Danecek, et al., 2016; Loh, Palamara, et al., 2016) as supplied by Eagle. As comparisons, we extracted 96 individuals from selected 1KGP populations to construct the recombination map. In the few cases where fewer than 96 individuals were available from a population in 1000 Genomes such as ASW, CDX, GBR, LWK, MSL, and MXL, we extracted the number of individuals based on the next available precomputed LDhat table. For CEU, we also downloaded OMNI array data (ftp://ftp.1000genomes.ebi.ac.uk/vol1/ftp/release/20130502) to help benchmark our pipeline. Next, each chromosome was split into windows of 4000 SNPs with 200 overlapping SNPs between windows. Finally, per previous studies (Auton et al., 2015; Xue et al., 2020; Zhou et al., 2020), the *interval* program from LDhat was run with the following settings: 30 million iterations, sampling set to every 15,000 iterations, 7.5 million iterations used for burn-in, and a block penalty of 5. The resulting estimates were integrated by removing the distal half of each overlapping region for each window and combining all windows for each chromosome.

The LDhat *interval* program estimates population-scaled recombination rate for a set of individuals using a composite likelihood method (Auton & McVean, 2007). The sex-averaged recombination rate, r, can be recovered by the relationship ρ = 4N_e_r, where N_e_ is the effective population size. Following previous protocol (Auton et al., 2015; Kong et al., 2010), we regressed the population-scaled estimates from LDhat for each population on the sex-averaged recombination rate from deCODE in order to estimate 4N_e_ (**Supplemental Table 1**). Regression was performed by aggregating corresponding rates in 5 Mb windows across the autosomes. After rescaling by the inferred 4N_e_ for each population, we then compute the recombination rate, r, per locus in centimorgan per megabase (cM/Mb).

To benchmark our pipeline, we applied it to OMNI array genotypes for 96 randomly selected CEU individuals from 1KGP and then compared the inferred map from our pipeline to the publicly available map (downloaded from ftp://ftp.1000genomes.ebi.ac.uk/vol1/ftp/technical/working/20130507_omni_recombination_r ates), which was originally also inferred from the OMNI array data.

We also inferred a recombination map for NH using IBDrecomb (Zhou et al., 2020), which uses the end points of IBD segments to infer recombination rates for a given set of individuals. IBDrecomb has been shown to perform similarly to LDhat while outperforming admixture-based methods on simulated and real data. To infer the recombination map using IBDrecomb, we first called IBD segments using Refined IBD (B. L. Browning & Browning, 2013b) (version 17Jan20.102) with default settings on genotypes phased using the omnibus map. We then run the *merge-ibd-segments* tool with 1 allowed error and maximum distance of 0.5 cM. Finally, we infer maps at 10 kb scale using these merged segments with the remaining parameters at default settings. Because a large number of IBD segments are needed to have sufficient number of informative historical recombination events, we used all individuals in our Native Hawaiian cohort to detect IBD segments before inferring a recombination map with IBDrecomb. The inferred maps using both LDhat and IBDrecomb are released on https://github.com/bldinh/recombination_maps in hg38.

### Evaluating the impact of population-specific recombination map on downstream statistical and population genetic applications

With each constructed Native Hawaiian-specific recombination map, we evaluated its impact on four distinct statistical and population genetic applications where information of the recombination landscapes are usually required for inference: imputation, local ancestry inference, IBD segment calling and relatedness inference, and genome scans of positive selection. In each case, we compared the performance and efficacy between using a Native Hawaiian-specific map (NH map) or the default omnibus map released by Eagle (Eagle map). We generally used the NH map for evaluation as most analyses incorporated the entirety of the admixed NH study samples that were not used in map constructions. In all cases, we excluded the 192 individuals that were used to construct both Native Hawaiian-specific maps from evaluation.

### Imputation

We phased and imputed our Native Hawaiian cohort (removing individuals that were used to construct the recombination map after phasing) using either the NH map or the omnibus Eagle map with HGDP+1KGP individuals (N = 3942) from gnomAD v3.1 as the imputation reference panel (Karczewski et al., 2020). Phasing was completed using Eagle v2.4.1 (Loh, Danecek, et al., 2016; Loh, Palamara, et al., 2016) with the same reference, and imputation was performed in-house using Minimac4 version 1.0.2 (Das et al., 2016; Fuchsberger et al., 2015; Howie et al., 2012). To evaluate the imputation accuracy, we compared the imputed genotype to exon sequencing data available for a subset of 607 MEC-NH individuals. Imputation accuracy was measured by the squared Pearson correlation of the sequenced genotype to the imputed dosage for variants that are found in both datasets. In total, we compare 1,248,599 genotypes across 607 individuals and 2057 SNPs.

### Local ancestry inference

We modeled Native Hawaiian individuals with four ancestry components corresponding to those found most prevalently in Africa, East Asia, Europe, and Polynesia (Lin et al., 2020). A total of 708, 800, and 671, individuals were selected from the HGDP+1KGP (from gnomAD v3.1) representing African, East Asian, and European ancestries (**Supplemental Table 2**). In addition, we added 176 MEC-NH individuals previously estimated to have more than 90% Polynesian ancestry to create a set of reference individuals for local ancestry inference (Lin et al., 2020; Sun et al., 2021) (after removing any individuals used in recombination map constructions). In total, 3,665 MEC-NH individuals not used as ancestry reference or in map construction were available to assess local ancestry inference (LAI) concordance. We pre-phased every MEC-NH individual in the study sample together with the HGDP+1KGP reference individual before separating the reference and test sets for LAI with RFMIX v2.03-r0 (Maples et al., 2013). Because recombination maps are also used during phasing, we performed our evaluation in two main comparisons: (1) comparison of RFMIX inference using the same initial pre-phased data using the NH map and (2) comparison of RFMIX inference after separately phasing the data on different maps. The former compares the direct effect a recombination map has on LAI and the latter encompasses the difference a recombination map would make on an analysis pipeline.

To compute concordance of LAI, at each site output by RFMix and for each individual we calculated the joint probabilities of the inferred ancestry between the recombination maps. We compared these inferences in a similar manner to a previous local ancestry study (S. R. Browning et al., 2016): if an individual has a segment inferred as belonging to ancestry A and ancestry B using map 1 and from ancestry C and ancestry A using map 2, we compare the inferred haplotypes so as to maximize their agreement. In this example, we calculate the inference as both maps agreeing on haplotype A and disagreeing for the other. Our joint probability calculation uses the marginal probabilities for this maximal haplotype combination.

### Integrated haplotype score

To evaluate the impact of recombination maps on genomic scans for signatures of positive natural selection, we randomly selected 150 individuals from our Native Hawaiian cohort and calculated iHS values using the selscan program (Szpiech & Hernandez, 2014). These scores were then normalized using the *norm* program included with selscan. We compared the normalized iHS results from our NH recombination map and two alternatives that would be commonly used in practice: the omnibus Eagle map and the pedigree-based European map from deCODE. We used a normalized z-score of |4| as the threshold for declaring a putative positively selected locus using either the omnibus or the pedigree map and examined the corresponding score on the NH map.

### IBD segment calling and inference of relatedness

Using genotypes phased previously for LAI (above), we called IBD segments using Refined IBD version 17Jan20.102 (B. L. Browning & Browning, 2013b). We then merged nearby IBD segments that could be spuriously broken up using the *merge-ibd-segments* tool provided by the authors of Refined IBD with the recommended parameters (1 allowed error, less than 0.6 cM distance). Initial analysis showed an elevated number of spurious IBD segments within both maps. For example, 70% of all detected IBD segments on chromosome 16 of length 15-20 cM map to the region located approximately at 15,000,000 to 17,000,000 bp. This region appears to have spuriously elevated recombination rate estimates; upon closer examination, the spurious pileup of IBD segments is due to genomic regions with very low variant coverage. We thus instituted a heuristic to create analysis masks (**below**) to be applied to the recombination maps before continuing IBD and relatedness analyses. We excluded the gap-filling step to reduce potential overestimations of shared IBD and excluded portions of IBD segments overlapping masked regions (additionally excluding them from the *length of genome* calculations that are specific to each map). We then used the IBD segments to infer genetic relatedness between any pair individuals by the following calculation: 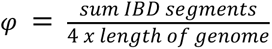, where *φ* is the kinship coefficient. We used KING (Manichaikul et al., 2010), an allele-frequency based method, to independently call genetic relatedness and restricted our comparisons due to the impact of different recombination maps to individuals estimated to be 3^rd^-degree relatives and higher (kinship estimate > 0.0442). Our comparisons were restricted to the intersection of pairs of individuals with estimated shared IBD segments on both maps.

### Analysis masks for recombination maps

Upon closer inspection of the distribution of IBD segments generated using the default omnibus map and the NH map, we found an increase of segments with length ranging between 15-30 cM. We noticed that genomic regions with sparse SNP density and the resulting elevated inferred rate of recombination would lead to accumulations of spurious IBD segments. These spurious accumulations of lengthy IBD segments had been previously noted (Ramstetter et al., 2017), and if unadjusted, will lead to erroneous inference of familial relationships. To avoid inaccurate results in downstream applications of the recombination maps, we created a filter to detect regions with conditions that may bias analyses. In tiling windows of 500 kb length (shifting in 50 kb increments), we flagged the region within the window to be considered for removal from downstream analysis if the number of markers available is less than 15 (compared to an average of 242.50 markers per 500 kb across the autosome for our Native Hawaiian cohort on the MEGA array). Other thresholds were considered, but we defined our threshold to attain a balance between detection of regions that are outliers and minimizing the total length of regions filtered out. Specifically, for our IBD analyses, we effectively set the genetic distance across these regions to 0 cM in order to dampen the inflation of inferred segment lengths and counts seen in our initial analysis. In total, our masks span 160.94 Mb (5.60%) of the autosome, and the coordinates for the Native Hawaiians on the MEGA array in hg38 can be found in **Supplemental Table 3**.

## Results

### Benchmarking the recombination map inference pipeline

We implemented a pipeline to generate recombination maps using LDhat (**Materials and Methods**). Notably, LDhat infers the relative, population-level recombination rate, ρ. Population-level rates, and the LD information used in inferring these rates, can be influenced by past population sizes. Therefore, following previous practices, we inferred and regressed out the contribution of the demographic history as measured by effective population size, N_e_ (**Materials and Methods**), in order to better compare the inferred recombination landscape across populations (but also see **Discussion** for limitations). We benchmarked our pipeline by applying it to OMNI array genotypes for 96 randomly selected CEU individuals from 1KGP and comparing the resulting map from our pipeline to the publicly available map originally also inferred from the OMNI array data. Overall, we observed high correlations (r = 0.93 to 0.96) across all scales of the map, ranging from 50 kb to 5 Mb (**Table 1**). The correlation was lower at the finest scale of 10 kb (r = 0.86), likely due to noise in LD estimates at fine scales given finite sample size (Bhérer et al., 2017). We also extracted the set of OMNI array SNPs from the public release of 1KGP phase 3 sequencing data for the CEU population to infer the recombination map, which again showed high correlation across all scales when compared to that inferred from the OMNI array genotype using the same in-house pipeline (r = 0.91 – 0.98; **Table 1**). Therefore, we concluded that our implemented pipeline near-faithfully recreated the previously published map, and that, given the same SNP content, differences in the distribution of any genotyping errors attributed to data generation platform (OMNI array vs. low pass sequencing) did not substantially impact the recombination map inference.

**Table 1:** Benchmarking LDhat inference pipeline. Each map was inferred from 96 randomly selected individuals from the corresponding 1KGP population via our LDhat pipeline. The inferred map would either be based on OMNI array genotypes, OMNI array SNPs extracted from WGS genotypes, or MEGA array SNPs extracted from WGS genotypes. The inferred maps were compared to the corresponding 1KGP recombination map for the same population or another inferred map using our in-house pipeline. We measured concordance between maps by the Pearson correlation coefficient. The first comparison established that our implemented pipeline faithfully recreated the published recombination map for CEU using the same input dataset. The second comparison established that the inferred map is robust to any differences in genotype quality between genotyping or sequencing. The third comparison established that maps based on the set of OMNI array SNPs or MEGA array SNPs are highly concordant, at least for 50kb scale or higher. The last comparison in YRI suggested that maps inferred using MEGA array genotypes can be compared directly to published 1KGP maps.

| Inferred Map Data Source |  |  | Comparison Data Source | Resolution |  |  |  |  |
| --- | --- | --- | --- | --- | --- | --- | --- | --- |
| Population | SNP Set | Data Source |  | 10kb | 50kb | 100kb | 1Mb | 5Mb |
| CEU | OMNI | Array | Published | 0.86 | 0.93 | 0.94 | 0.95 | 0.96 |
| CEU | OMNI | WGS | OMNI array data, in-house | 0.91 | 0.95 | 0.95 | 0.96 | 0.98 |
| CEU | MEGA | WGS | OMNI SNPs from WGS data, in-house | 0.78 | 0.91 | 0.93 | 0.94 | 0.97 |
| YRI | MEGA | WGS | Published | 0.82 | 0.92 | 0.94 | 0.95 | 0.97 |

Because the Native Hawaiian genetic data from the MEC is only available on the MEGA array, we also benchmarked the impact using a different set of SNPs on map inference. Again, using the CEU 1KGP population as a model, we extracted the genotypes found on the MEGA array to infer a recombination map. The resulting map is highly correlated with the version generated based on extracted OMNI array SNPs, at least for 50 kb scale or higher (r > 0.91; **Table 1**). Taken together, our LDhat inference pipeline, when applied to SNPs found on the MEGA array for the CEU population, would produce highly concordant recombination maps at 50 kb scale or higher compared to the published map. Therefore, the recombination landscape for Native Hawaiians that we inferred here can be compared to those published for other 1KGP populations. Nevertheless, for between-population comparisons of the recombination landscape we compared the inferred NH maps to those based on the MEGA array SNP content extracted from 1KGP sequence data.

### Contrasting the Native Hawaiian recombination landscape

We proceeded to infer a recombination map for 96 randomly selected admixed Native Hawaiians from our study sample (NH map) to represent the current Native Hawaiian population and a map for 96 individuals from our study sample previously estimated to be most enriched with the Polynesian ancestries found predominantly in Native Hawaiians (PNS map; **Materials and Methods**). The PNS map was constructed to provide insights on the recombination landscape of the ancient Polynesians and how it may differ with other continental populations around the world. However, we caution that the Polynesian ancestries found predominantly in Native Hawaiians may or may not have diverged from those found in other Polynesian populations such as the Samoans. Consistent with their known history of serial bottleneck and long-term isolation (Chiang, 2021), the inferred past population size, parameterized by the effective population size (N_e_), was substantially smaller for PNS than other populations in 1KGP (**Supplemental Table 1**). The estimated N_e_ for NH was larger, more in line with other continental and admixed populations in 1KGP. Regressing out the effect of demographic history (**Materials and Methods**), we then compared the inferred Native Hawaiian recombination landscape with that from other populations that were similarly constructed using MEGA array genotypes extracted from the 1KGP phase 3 datasets.

We found that populations with recent shared history displayed higher levels of correlation, as expected (**Figure 1a**). This is notable for populations representing each of the ancestral components found: African (MEC-AA, ASW, and YRI), East Asian (CHB, CHS, and JPT), and European (CEU, TSI). At the 50 kb scale, the recombination landscape as indicated by the NH map showed intermediate correlations with the non-Native Hawaiian populations (r = 0.865 - 0.899), tending to correlate best with populations representing East Asian (r = 0.892 – 0.898) and European ancestry (r = 0.895 - 0.899), likely due to substantial admixture from these two ancestral components. In contrast, the landscape from PNS showed the lowest correlations with almost all populations compared (r = 0.774 - 0.847). PNS correlated most strongly with NH (r = 0.847) and most weakly with Peruvian in Lima, Peru (PEL; r = 0.774). Overall, PNS correlated more poorly with other populations than PEL, a population with a known history of isolation. Taken together, our results are consistent with PNS representing a previously unrepresented component of ancestry by 1KGP and may suggest a stronger effect of isolation or enhanced genetic drift in the past compared to PEL (although we also note that we did not prune PEL to enrich for Indigenous American ancestry due to small sample size).

**Figure 1:**
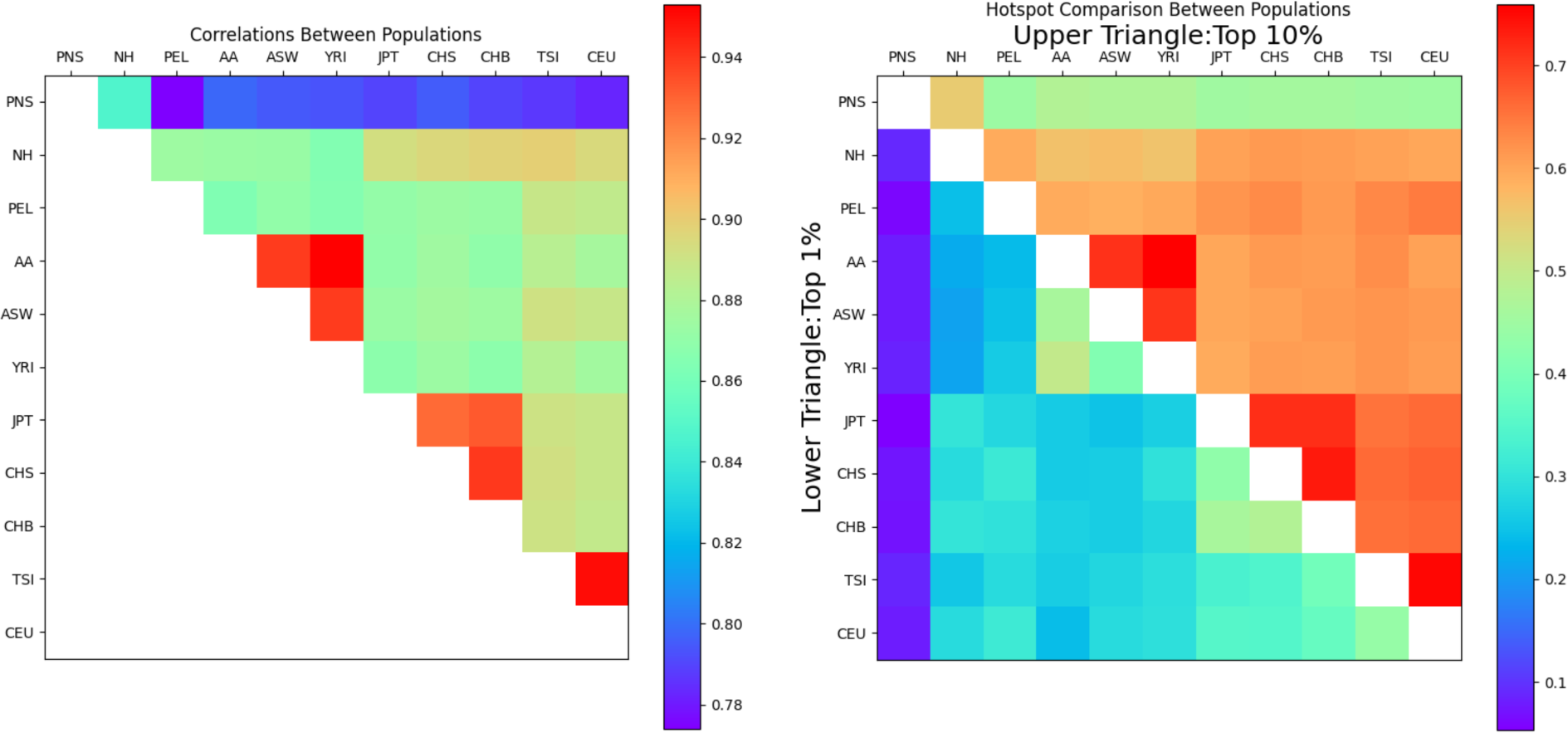
Correlation of recombination landscape and hotspots between populations. Correlations between populations at 50 kb scale (**A**) and sharing of recombination hotspots defined as the top 1% (lower left) or 10% (upper right) windows 50 kb in size by inferred recombination rate (**B**). Both the correlations and proportion of sharing were calculated by excluding genomic regions that fall within the analysis masks (**Materials and Methods**). All landscape and hotspot comparisons here were based on recombination maps inferred in-house using MEGA array SNPs from genotyping (PNS, NH, AA) or extracted from WGS data (1KGP populations with Indigenous American-[PEL], African-[ASW, YRI], East Asian-[JPT, CHS, CHB], and European-ancestries [TSI, CEU]). AA is African American data from MEC.

These trends were also observed when we compared recombination hotspot sharing between populations (**Figure 1b**). Following methodology of previous studies (Wegmann et al., 2011), we divided the genome into 50 kb windows and defined hotspots for each population as the windows in the top 1 percentile or 10 percentile with the highest average recombination rate. For each pair of populations, we then compared the shared proportion of hotspots. The relationships at both hotspot thresholds were similar to the ones observed for correlations of the genome-wide recombination landscape (**Figure 1a**): populations expected to represent recent shared history had the highest sharing at both thresholds. In contrast, PNS had lower hotspot sharing relative to other populations except for NH, with the lowest sharing with PEL. Compared to PNS, the NH map has better relative sharing with the other 1KGP populations, including slightly higher sharing with European and East Asian populations. PNS’s low correlation genome-wide in recombination rate variations and hotspot sharing with other populations is thus indicative of a unique recombination landscape.

### Impact of the recombination map on imputation

We evaluated the impact of a population-specific map for Native Hawaiians on downstream statistical and population genetic applications, specifically focusing on the impact of the recombination map on genotype imputation, local ancestry inference, IBD and relatedness inference, and genomic scans for positive selection.

Imputation is a common analysis that uses haplotype sharing to infer genotypes at variants not directly observed in a given dataset. The recombination map is needed for phasing (a pre-requisite for imputation) and for imputation itself as it provides the prior that the ancestral haplotype may have switched from one haplotype to another at a given genomic location. We thus evaluated the impact of using a potentially mis-specified omnibus map (*i.e.* the default recombination map released by the commonly used phasing software, Eagle (Loh, Danecek, et al., 2016; Loh, Palamara, et al., 2016)) in phasing and imputation, compared to the population-specific map constructed in the present study.

We imputed 453 and 154 Native Hawaiian individuals genotyped on the Illumina Human660W (NHBC) and MEGA arrays, respectively, against the 1KGP+HGDP reference panel released by gnomAD (Karczewski et al., 2020). Compared to the targeted exon sequencing data we have on overlapping individuals, we were able to evaluate imputation accuracy in the 658 and 482 SNPs for NHBC and MEGA cohorts, respectively. Across three minor allele frequency (MAF) bins, 0.5 – 1%, 1 – 5%, and 5 – 50%, we observed no notable difference in imputation accuracy when imputation was performed using the NH map compared to the omnibus map (**Figure 2**, **Supplemental Table 4**)

**Figure 2:**
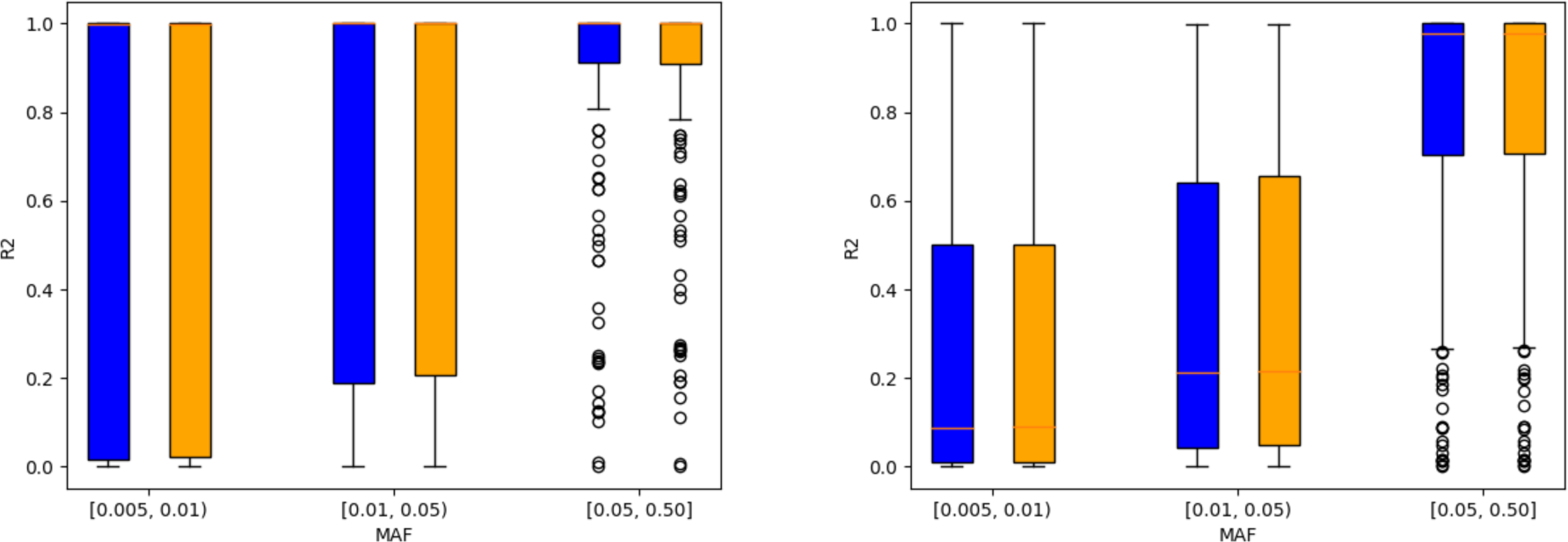
evaluation of the impact of recombination maps on imputation accuracy. Comparison of imputation accuracy using the NH map (blue) and the omnibus map (orange). MAF for each SNP was computed using the sequencing data. **Left**: 154 individuals on the MEGA array were compared on 482 available SNPs. Restricting to MAF ≥0.5% resulted in bins with 134, 112, and 168 SNPs, respectively. **Right**: 453 individuals on the Human660W array were compared on 658 overlapping SNPs. More SNPs are available for comparison here due to larger number of individuals available for analysis. Restricting to MAF ≥0.5% resulted in bins with 64, 94, and 165 SNPs, respectively.

### Impact of recombination map on local ancestry inference

We inferred the local ancestries across 3,665 Native Hawaiian individuals genotyped on the MEGA array using RFMIX (Maples et al., 2013), with either the NH map or the omnibus map that provided information of recombination rates genome-wide. Overall, there is a large agreement of local ancestries inferred based on the NH map and those based on the omnibus map (concordance = 97.2%, **Table 2**), particularly when phasing was performed only once before local ancestry inference with two separate maps. If we separately phased and ran RFMIX with different maps, the resulting concordance lowered to 95.8% (**Table 2**), suggesting that the choice of the recombination map could make a difference in the LAI calls, though perhaps not strongly. These results are also consistent with previous findings that even a naïve uniform recombination map would produce <10% of discordance in inferred ancestry calls (Sun et al., 2021).

**Table 2:**
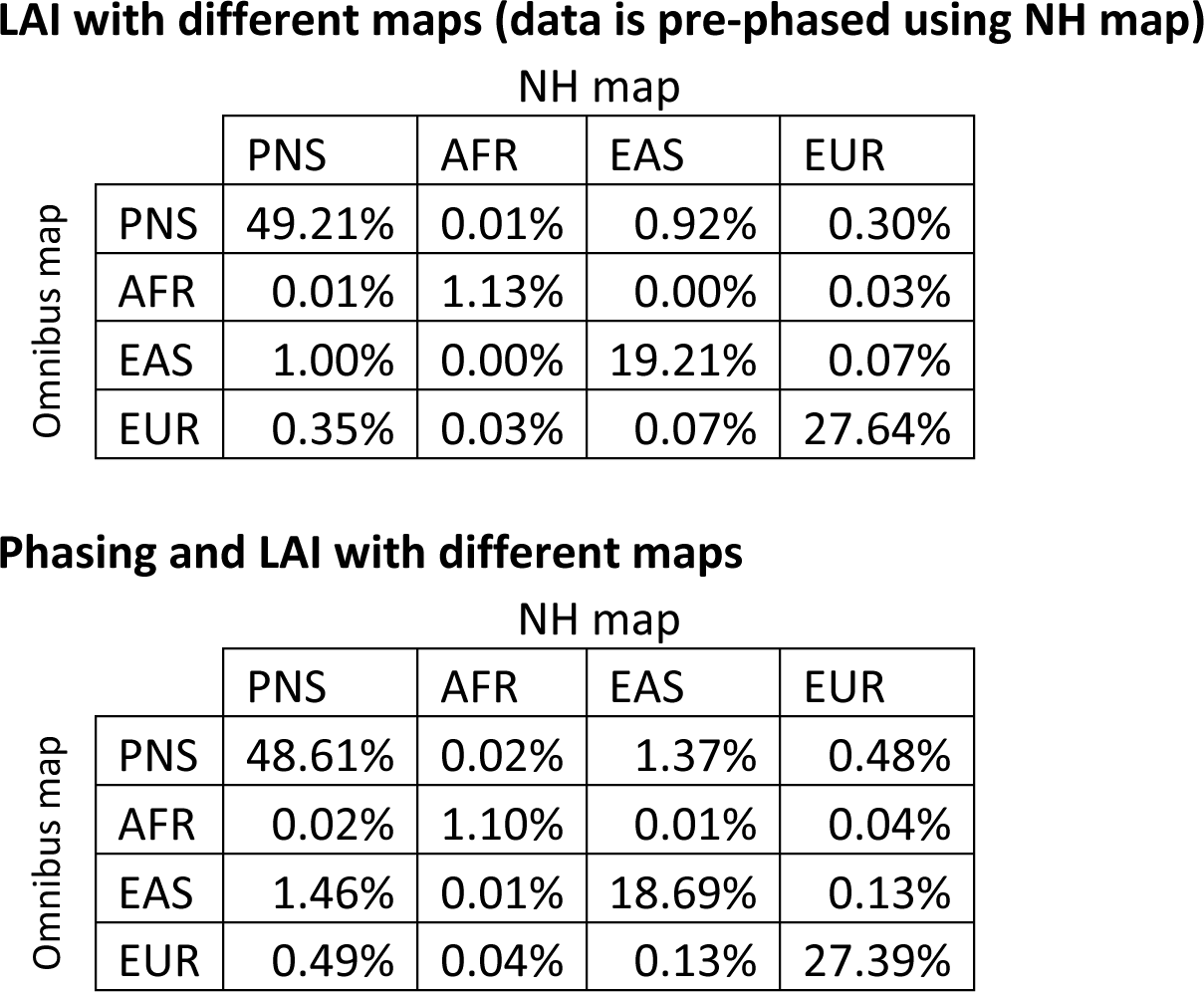
Concordance of inferred local ancestry calls based on different recombination maps. The Native Hawaiian population is modeled as a four-way admixture, with ancestry components from Polynesia (PNS), Africa (AFR), East Asia (EAS), and Europe (EUR).

|  |  | NH map |  |  |  |
| --- | --- | --- | --- | --- | --- |
|  |  | PNS | AFR | EAS | EUR |
| Omnibus map | PNS | 49.21% | 0.01% | 0.92% | 0.30% |
|  | AFR | 0.01% | 1.13% | 0.00% | 0.03% |
|  | EAS | 1.00% | 0.00% | 19.21% | 0.07% |
|  | EUR | 0.35% | 0.03% | 0.07% | 27.64% |

**Phasing and LAI with different maps**
|  |  | NH map |  |  |  |
| --- | --- | --- | --- | --- | --- |
|  |  | PNS | AFR | EAS | EUR |
| Omnibus map | PNS | 48.61% | 0.02% | 1.37% | 0.48% |
|  | AFR | 0.02% | 1.10% | 0.01% | 0.04% |
|  | EAS | 1.46% | 0.01% | 18.69% | 0.13% |
|  | EUR | 0.49% | 0.04% | 0.13% | 27.39% |

### Impact of recombination map on IBD segment calling and relatedness inference

Our initial IBD analysis identified regions of the genome where we observed extreme pileups of IBD segments that are spurious (>90% of individuals would share IBD segments locally). We thus created analysis masks to remove these spurious regions driven by insufficient SNP coverage (see **Materials and Methods**, **Supplemental Figure 1**). When comparing the distribution of IBD segments using masked versions of both maps, we observed little difference (**Supplemental Figure 1**). Given that the inferred recombination rates in regions of sparse SNP density could be unreliable and confound downstream analysis such as IBD segment calling, we also released a set of masked regions (**Supplemental Table 3**) for users to apply when using these recombination maps.

We used the kinship coefficient (*φ*) based on the pairwise proportion of IBD sharing to compare differences in relatedness inference due to the recombination map (**Materials and Methods**). Of the 218,132 pairs of individuals inferred by KING to be 3^rd^-degree or closer relatives, 200,617 were found in both the omnibus and NH maps to compare. The estimated kinship coefficient from each map was calculated independently and the maps were found to be highly concordant with a Pearson correlation coefficient of 0.995.

### Impact of recombination map on genomic scans for positive selection using integrated haplotype score

Lastly, we evaluated the impact of recombination maps on the robustness of a haplotype-based metric of positive selection, the integrated haplotype score (iHS) (Voight et al., 2006). We considered loci with normalized iHS absolute Z-score (|Z|) > 4 as candidate loci under positive selection. Most SNPs in our scans using different recombination maps were concordant with similar iHS values (**Figure 3**). However, we identified 6 SNPs across 4 loci (nearby genes: DISP1, TLCD1, FAM222B, and WNT7B) that showed |Z| values > 4 when using the omnibus map but normalized |Z| values between 0 to 2 when using the NH map (**Supplemental Figures 2-5**). Moreover, there are additional loci that would have attenuated evidence of selection in analysis using the NH map (|Z|< 2) but fall short of our conservative cutoff of 4 standard deviations when using the omnibus map. Selection scans may also be performed using a pedigree-based map, such as one from deCODE, to avoid any confounding between haplotype-based statistics and local LD patterns. We also compared the analysis using the deCODE pedigree map (**Supplemental Figure 6**), which is based on a European-ancestry population in Iceland. We observed 46 outlier SNPs across 19 loci that would have attenuated evidence of selection (|Z| > 4 using deCODE map, but < 2 using NH map). Due to the substantial increase of candidate loci under positive selection with the deCODE map and the overall larger iHS after normalization (the largest |Z| near 15), it is clear that haplotype-based selection analyses are sensitive to the choice of recombination maps. Therefore, using the population-specific recombination map in haplotype-based selection scans may better protect against spurious detections of loci under positive selection.

**Figure 3:**
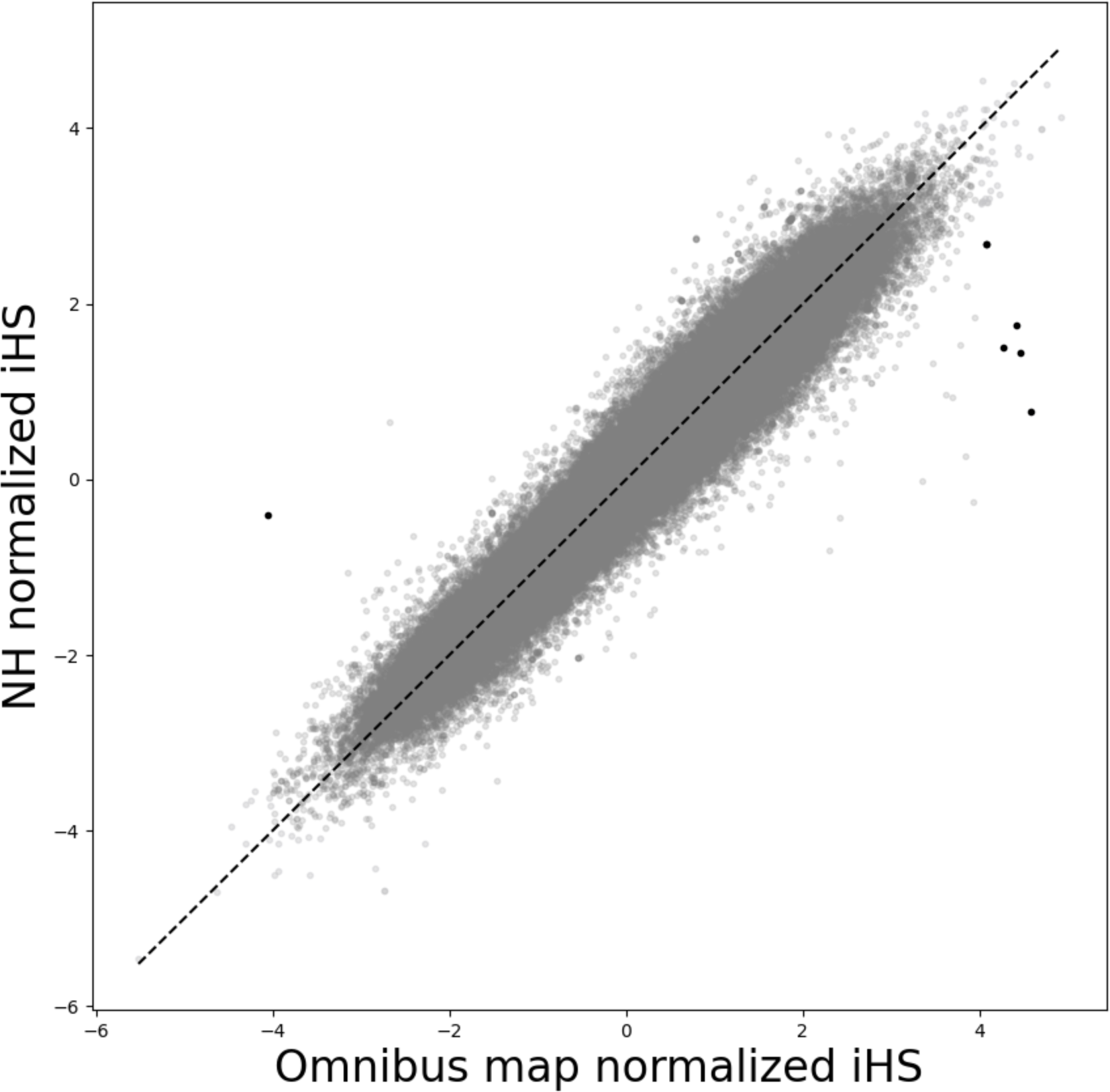
Genomic scan for positive selection using haplotype-based statistics is sensitive to choice of recombination map. Genomic scan of adaptation was measured using iHS as a metric, as implemented in selscan. We estimated iHS for 486,600 SNPs in 150 Native Hawaiian individuals from the MEC genotyped on the MEGA array using either the omnibus map (x-axis) or the NH LDhat (y-axis) map. After normalization by selscan, a score of |z| > 4 is considered genome-wide significant. We identified six SNPs across four loci that would have significant evidence of adaptation using iHS based on the omnibus map, when all of the statistical evidence would be much more attenuated if a population-specific map was used.

### A recombination map based on IBD segments

In addition to LDhat, an alternative approach to infer genome-wide recombination maps based on array data is to utilize information from IBD segments. As larger sample size is needed to have sufficient number of IBD segments for inference, we inferred a recombination map using 3,937 individuals using IBDrecomb (Zhou et al., 2020), which would be more akin to the NH map constructed using LDhat. The resulting IBD-based map showed large concordance with the NH map (**Supplemental Table 5**), although the correlation may be lower compared to that reported previously for African Americans (Zhou et al., 2020). For instance, the correlation between an IBD-based map of an African American cohort and an LDhat map from ASW was previously reported to be as high as 0.95 at the 500kb scale; we observed a correlation of 0.90, 0.90, and 0.92 at the 500 kb, 1 Mb, and 5 Mb scales (**Supplemental Table 5**). This may be due to the differences in the precision in IBD segment detection using array versus sequencing data (B. L. Browning & Browning, 2013a; Chiang et al., 2016), which may percolate to the precision in the constructed recombination maps, or it may be due to differences in ancestry composition between our African American cohort (sampled largely in Southern California) with that tested previously (Zhou et al., 2020). In comparison, the correlation between IBD-based map and LDhat map from NH are between 0.85 to 0.89 across the same scales (**Supplemental Table 5**). Nevertheless, we found that compared to the omnibus map, the IBD-based map also did not substantially impact the downstream inference for imputation (**Supplemental Figure 7**), LAI (**Supplemental Table 6**; 94.9% concordance, comparing IBD-based to LDhat-based map), and kinship coefficient estimates (Pearson correlation with NH map: 0.9945). We also found that genomic scan for positive selection using iHS would be more robust if we were to use an IBD-based recombination map for NH as well (**Supplemental Figure 8**), confirming our previous observation that a population-specific recombination map would be important for performing and interpreting haplotype-based analyses.

## Discussion

In the present study we implemented a pipeline to infer recombination maps for populations using LDhat. By utilizing both array and sequence data from 1KGP populations and comparing to the corresponding publicly available recombination maps, we demonstrated that our implemented pipeline could faithfully produce precise recombination maps. We also showed that recombination maps based on SNP content on the MEGA array have high concordance with published 1KGP maps. These results suggest that our constructed maps for the Native Hawaiian population based on the MEGA array data would be comparable to those already available for 1KGP populations based on the OMNI array.

To characterize the recombination landscape of Native Hawaiians and to generate a commonly used genomic resource for future genetic studies in Polynesian-ancestry individuals, we created two recombination maps, the PNS map and the NH map, from Native Hawaiian individuals estimated to have the highest proportion of Polynesian ancestry and individuals randomly sampled from the cohort, respectively. The former map was constructed to provide insights on the recombination landscape in the ancestral Polynesians prior to European colonization and subsequent waves of immigrations. However, we note that Polynesian ancestries are complex. This component of ancestry is one that is found prevalently in the MEC Native Hawaiian cohort, but could represent a mixture of Polynesian, Austronesian, and other ancestries that are unique to our sample. Therefore, a future evaluation of the applicability of the PNS map to other Polynesian populations across the Pacific would be of interest. The admixture history specific to the Native Hawaiians also contributed to the health disparity experienced by the population today, and the genetic risks to diseases must be evaluated considering this history (Sun et al., 2021; Taparra et al., 2021). As such, we also constructed the latter map, to better reflect the genomic composition of the Native Hawaiians today. The NH map would be more suited for genetic epidemiologic studies that use a population-based sample and is the primary map that we evaluated throughout this study.

Overall, we found that the recombination landscape from the PNS map stood out to be the most unique compared to other 1KGP and MEC populations based on the correlation of recombination rate variations across the genome and proportion of hotspot sharing. Notably, the pairwise sharing between YRI and CEU, two populations known to have divergent population histories, have higher correlation with one another (r = 0.876) than either population with PNS (0.793 and 0.783 for YRI and CEU, respectively). The landscape based on the NH map, in contrast, showed higher correlations with the maps from other populations, with slightly higher correlation with the maps inferred from European and East Asian populations. This is likely due to the recent admixture from other continental populations seen within the NH cohort today (Sun et al., 2021). Lastly, the PNS map showed the lowest correlation with PEL, and both PNS and PEL maps showed relatively low correlations with maps from other populations examined here. The Peruvian from Lima, Peru, is known for its history of isolation relative to other 1KGP populations (Harris et al., 2018). The observation here thus suggests that long-term isolation and lower effective population sizes are driving the LD pattern and shaping the recombination landscape. The low correlation between PNS and PEL, and between the two with other populations, are potentially reflecting their respective independent isolation histories.

We evaluated the impact of a population-specific map on downstream population and statistical genetic analyses. In general, we found little difference due to the choice of the recombination maps, particularly for imputation accuracy, local ancestry inference, and IBD segment detections and genetic relatedness inference. In local ancestry inference, we did observe an increased discordance between the two maps, if the input data were also phased separately using different maps (**Table 2**), suggesting that the choice of recombination maps would have a small but tangible impact on these downstream analyses. However, in each of these applications, the recombination maps are used mostly as priors while the actual inference is ultimately driven by the genetic data. Therefore, even though there are fine-scale differences in the recombination landscape between Native Hawaiians and other 1KGP populations, the accuracy of these downstream applications will not be substantially impacted by the choice of the map used.

Our findings suggest that the choice of the recombination map could make qualitative differences in population genetic analyses such as genomic scans for positive selection using the integrated haplotype score (iHS) because these statistics would incorporate recombination information in their calculation, representative of haplotype length surrounding a SNP. We found that 6 SNPs across 4 loci appeared to be putative selection loci (using a threshold of |iHS| > 4 after normalization) but have attenuated signals using the NH map (|iHS| < 2). The signals in the four loci are driven by the genes DISP1, TLCD1, FAM222B, and WNT7B. We examined each locus manually and did not find clear evidence for them to be under selection. DISP1 impacts signaling and transport for cellular proliferation and differentiation (Ehring et al., 2021). Mutations in the gene are associated with neuronal and endocrine diseases. TLCD1 affects membrane assembly and regulates membrane composition and tolerance to fatty acids (Ruiz et al., 2018). FAM222B has been associated with red blood cell regulation and protein binding using gene ontology (Luck et al., 2020). WNT7B is part of a pathway for embryonic developmental processes and is linked to carcinogenesis and cancer development, in particular gastric cancer (Gao et al., 2021; Goessling et al., 2009; Kirikoshi et al., 2001). Additional genes near the four loci can be found in **Supplemental Figures 2-5**. Overall, we found no suggestive functional basis to implicate these loci as under selection in Native Hawaiians.

We postulate that even though the NH map displayed elevated correlation of recombination rates genome-wide with many of the populations in 1KGP due to its recent admixture, it still accurately reflects the recombination events pertaining to the Polynesian ancestries as evidenced by its high correlation and overlap of hotspots with the PNS map. Assuming that much of the signals of positive selection should predate the recent admixture times over the last 10 generations or so, a population-specific map like the NH map captured much of the recombination landscape from the Polynesian ancestry component and should improve the robustness of haplotype-based scans of selection such as iHS. Indigenous populations such as the Native Hawaiians have been understudied with respect to their present-day medical conditions and healthcare considerations (Taparra, 2021). Their evolutionary or adaptive histories for surviving the most geographically isolated habitats on the planet for centuries could also contribute to the genetic causes of differences in disease risk between the Native Hawaiians and other continental populations. However, studies of indigenous populations have also been fraught with “just-so” stories that only superficially tied evolutionary hypotheses to apparent disparity in diseases (Chiang, 2021; Fox et al., 2020; Gosling et al., 2015). The availability of a population-specific recombination maps could help weed out false positive findings, thus making the resulting conclusions more robust and less harmful in future studies.

Finally, there are a number of limitations to our study, largely driven by the scarcity of genomic knowledge and data for Native Hawaiians. We chose to use LDhat to construct the recombination maps here as the available genomic data for Native Hawaiians are largely restricted to the MEGA array data. When WGS data become available for Native Hawaiians at scale, we would be able to evaluate more deeply the statistical genetic applications presented here. For instance, WGS data would allow us to evaluate the impact of recombination maps on imputation more thoroughly across the genome and different functional annotation. Additionally, WGS data would also allow us to model the demographic history of the Native Hawaiians, thus enabling the creation of more accurate recombination maps using methods, such as pyrho (Spence & Song, 2019a). On top of the improved accuracy by incorporating demographic history, pyrho can also infer recombination maps using hundreds of individuals, allowing us to reduce noise by including many more individuals. Many of the evaluations we have conducted here would be worth revisiting when this map based on sequenced data become available. Furthermore, LDhat infers the relative, population-scaled, recombination rates, ρ, rather than absolute recombination rates, r. Despite the effort to control for the impact of population history through estimating and regressing out the long-term effective population size, we presumed the differences in recombination landscapes observed across populations largely reflected differences in LD pattern due to the Hawaiian’s unique demographic history. We were unable to assess differences in recombination between Native Hawaiians and other populations due to, for example, PRDM9 motif usage variations (Hinch et al., 2011). To directly and reliably estimate the underlying recombination rates would require large-scale pedigree data, such as the one amassed by deCODE (Kong et al., 2010), but is extremely difficult to ascertain for a vulnerable, indigenous population such as Native Hawaiians. Nevertheless, the rates estimated by LDhat within Native Hawaiians are highly correlated with IBD-based maps, and these maps can inform and improve genetic analyses using haplotype-based statistics for this population, as we have evaluated here. This is thus an important step towards inclusion and lessening further irresponsible construct of selection stories for the Native Hawaiians. Finally, we also acknowledge that the underlying genetics and genomics should not supplant current standards through self-identity or genealogical records for defining community memberships; there is one Native Hawaiian population that cannot be discretized through genomics. The implications of these and future genetic findings on Pacific Islander health must be viewed through the lens of the social determinants of health with the goals to improve inclusion and equitable benefit sharing with the indigenous communities (Fox, 2020; Pineda et al., 2023).

## Acknowledgements

We would like to thank John Novembre and Vagheesh Narasimhan for discussions and comments on the preprinted version of this manuscript. We would also like to thank the Native Hawaiian participants in the Multiethnic Cohort that are involved in this study. The Multiethnic Cohort was funded through grants from the National Cancer Institute (U01CA164973, P01CA168530) and National Human Genome Research Institute (U01HG007397). We also would like to thank University of Hawai‘i Cancer Center’s Native Hawaiian Community Advisory Board for reviewing the study proposal and providing comments to earlier versions of this manuscript. This study is supported by grants from the National Institute of General Medical Sciences (R35GM142783 to C.W.K.C.) and the National Human Genome Research Institute (F31HG012159 to B.L.D.). Computation for this work is supported by the University of Southern California’s Center for High-Performance Computing (https://hpcc.usc.edu).

