## Supplemental Figures for "Recombination map tailored to Native Hawaiians improves robustness of genomic scans for positive selection"

**S1 Figure. Distribution of detected IBD segments with and without the analysis mask.** For the omnibus Eagle map and the NH LDhat map, we detected IBD segments using Refined IBD. Without applying the analysis mask, we noticed spurious accumulation of long IBD segments > 15cM (inset), driven by regions with low SNP density. These spurious segments are removed when the analysis mask was applied.

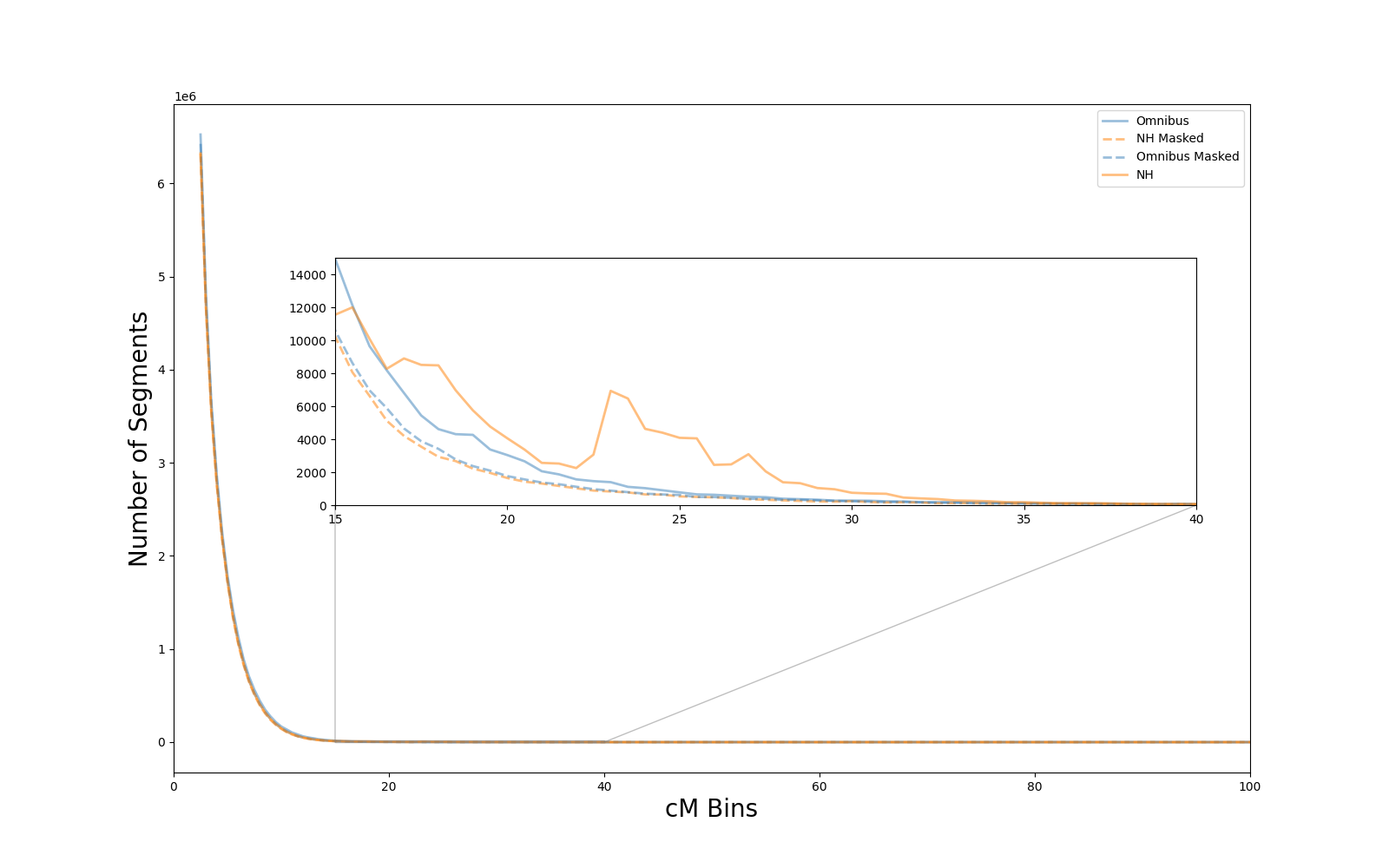

**S2 Figure. Likely spurious signal of adaptation on chromosome 1 detected by genomic scan using iHS with an omnibus map.** The largest iHS in this region occurs at position 222891307. In addition to signaling and transport for cellular proliferation and differentiation, DISP1 mutations and deficiencies are associated with neuronal and endocrine diseases in fetal development and cholesterol.

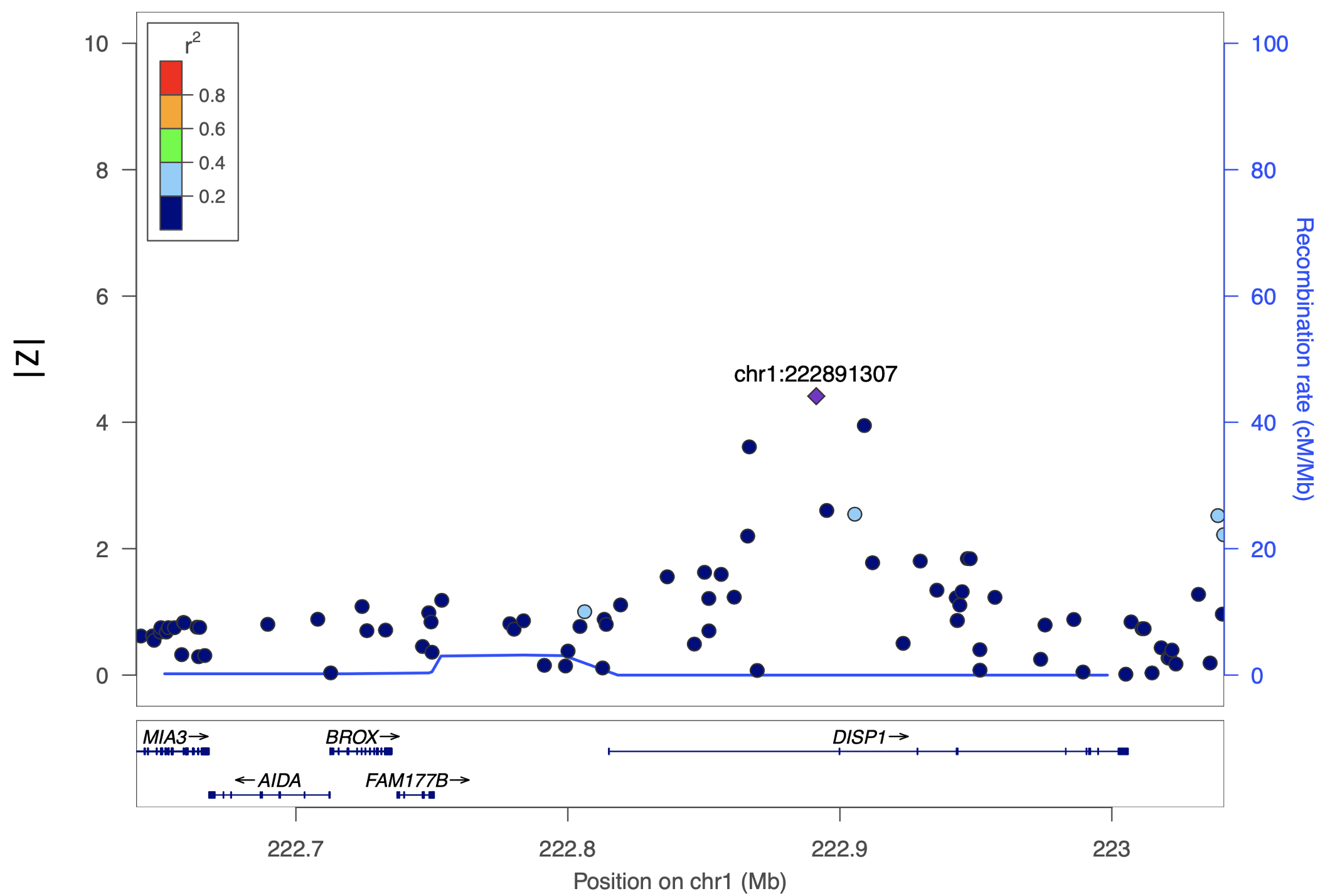

**S3 Figure. Likely spurious signal of adaptation on chromosome 6 detected by genomic scan using iHS with an omnibus map.** The largest iHS in this region occurs at 167726632. LOC441178 has been associated with oropharyngeal squamous cell carcinoma.

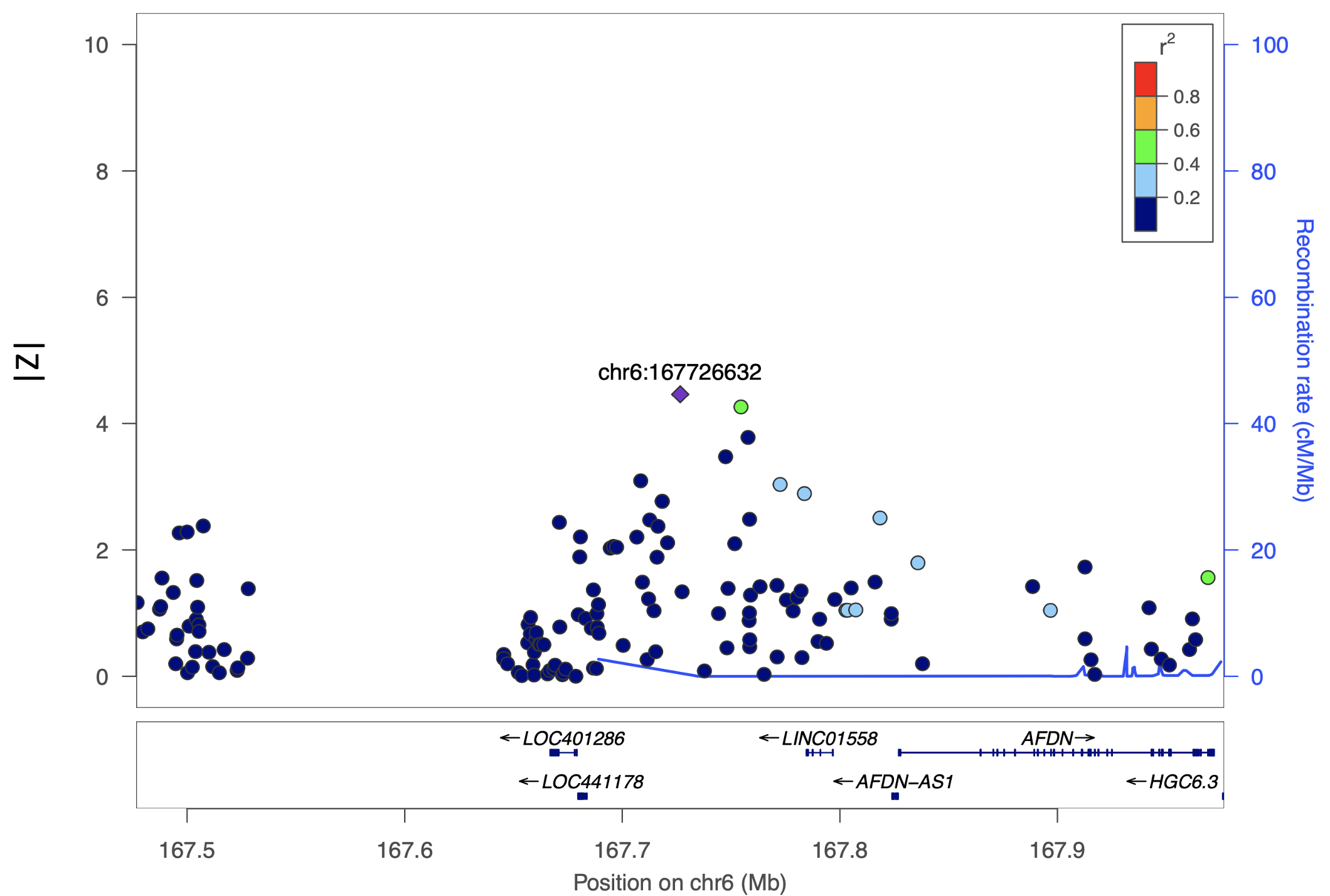

**S4 Figure. Likely spurious signal of adaptation on chromosome 17 detected by genomic scan using iHS with an omnibus map.** The largest iHS in this region occurs at position 28768467. In addition to red blood cell regulation and protein binding, FAM222B is also associated with blood protein measurement. TLCD1 inhibits incorporation of unsaturated fatty acids in membrane.

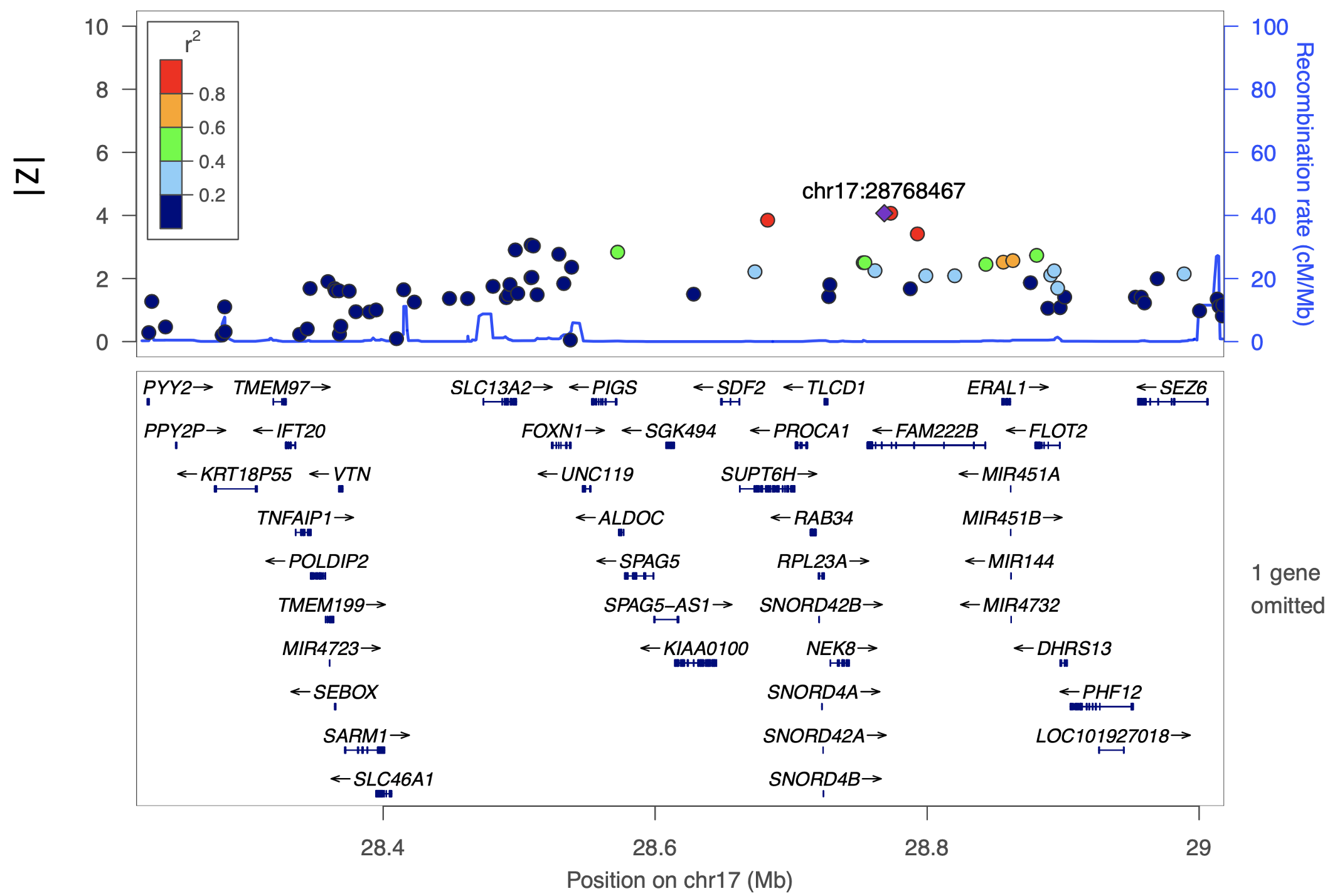

**S5 Figure.** **Likely spurious signal of adaptation on chromosome 22 detected by genomic scan using iHS with an omnibus map.** The largest iHS in this region occurs at position 45917335. In addition to the associations with various cancers, WNT7B is largely associated with gastric cancer.

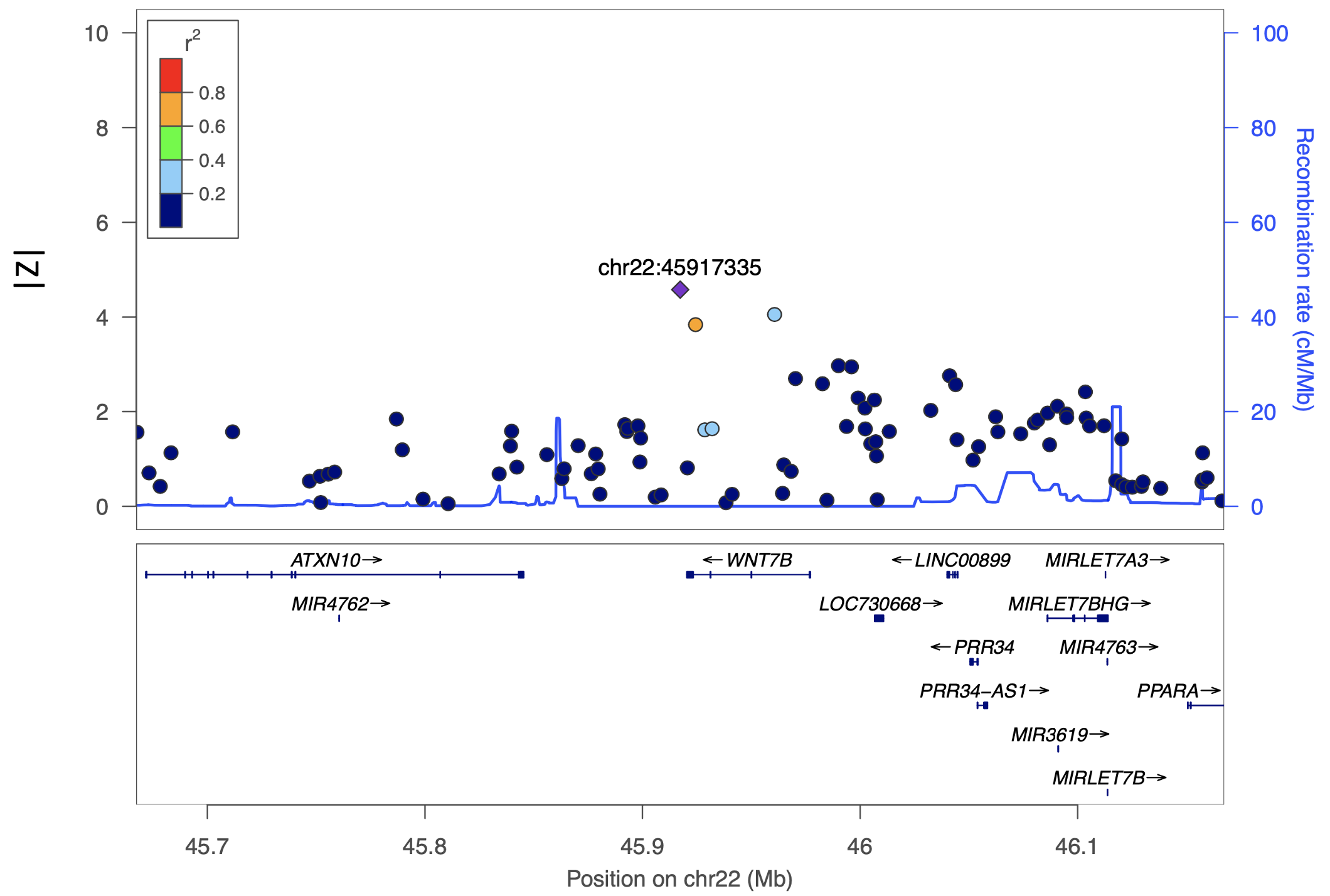

**S6 Figure. Comparison of iHS values using the deCODE map vs. NH map.** iHS scores were estimated for 150 MEC-NH individuals genotyped on the MEGA array. After normalization by selscan, a score of |z| > 4 was considered genome-wide significant. Black circles (46 total) indicate iHS found to be significant in deCODE analysis but not in NH map analysis. The large number of additional candidates from analysis with the deCODE map suggest the NH map is better suited for this selection scans for the Native Hawaiian population.

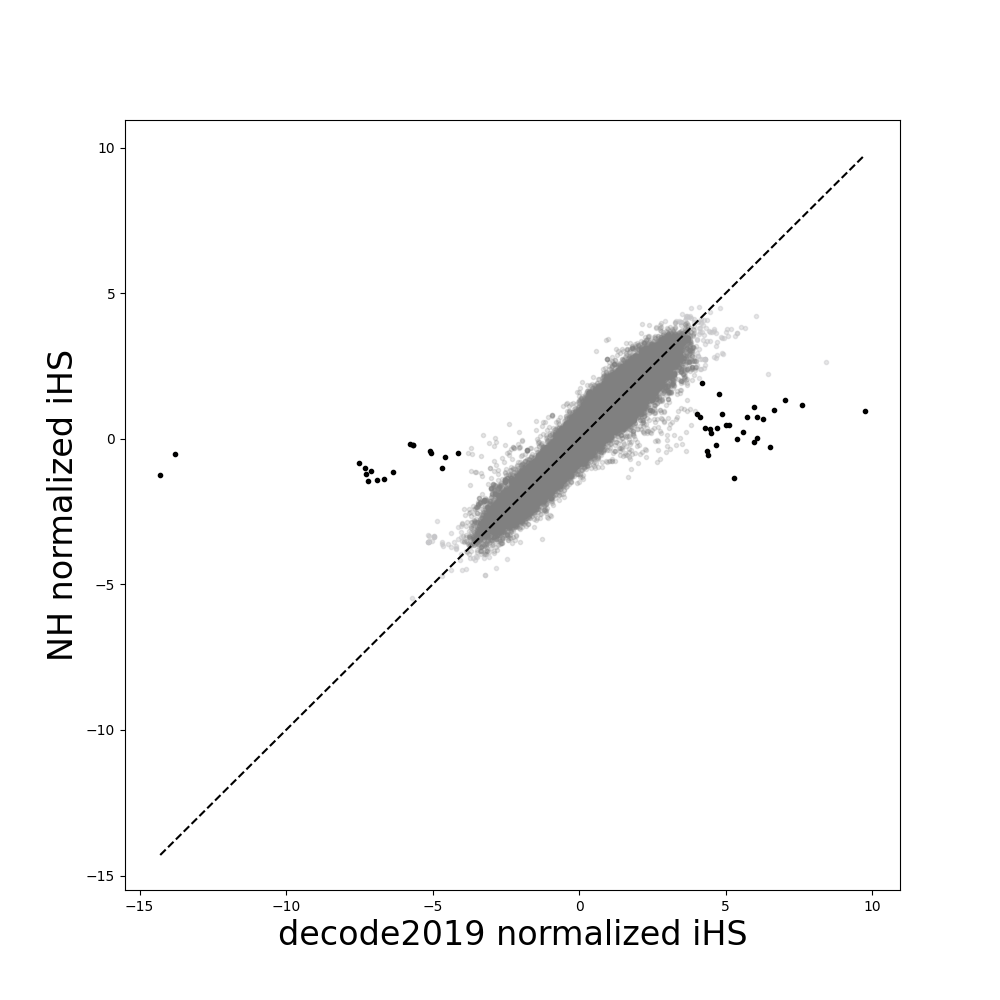

**S7 Figure.** **Evaluation of the impact of IBDrecomb map on imputation accuracy**

Comparison of imputation accuracy using the NH LDhat map (blue), the omnibus map (orange), and the NH IBDrecomb map (brown). MAF for each SNP was computed using the sequencing data. **Left**: 154 individuals on the MEGA array were compared on 482 available SNPs. Restricting to MAF ≥ 0.5% resulted in bins with 135, 112, and 168 SNPs, respectively. **Right**: 453 individuals on the Human660W array were compared on 658 overlapping SNPs. More SNPs are available for comparison here due to larger number of individuals available for analysis. Restricting to MAF ≥ 0.5% resulted in bins with 64, 94, and 165 SNPs, respectively.

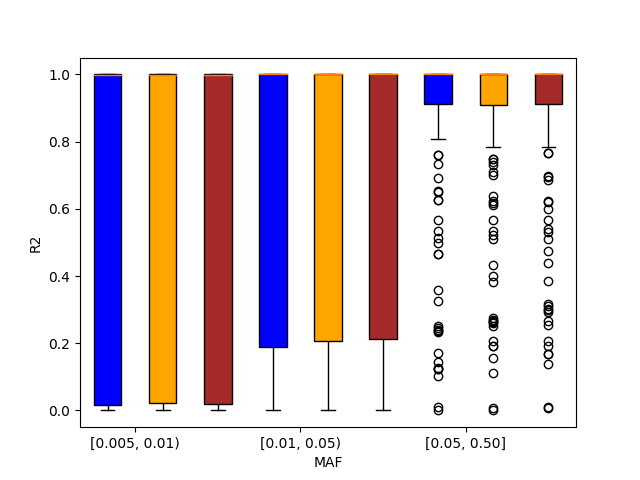

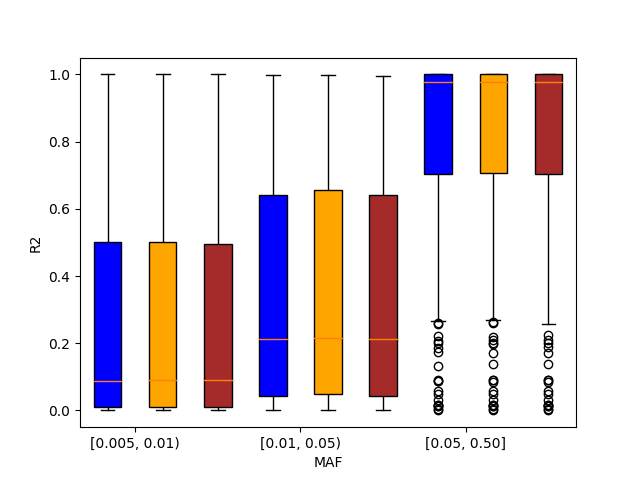

**S8 Figure. Comparison of iHS values using the omnibus Eagle map vs. NH IBDrecomb map.** iHS scores were estimated for 150 MEC-NH individuals genotyped on the MEGA array. After normalization by selscan, a score of |z| > 4 was considered genome-wide significant. iHS estimated using the NH IBDrecomb map are attenuated in comparison to the omnibus map (also seen in the comparison of the NH LDhat map to the omnibus map).

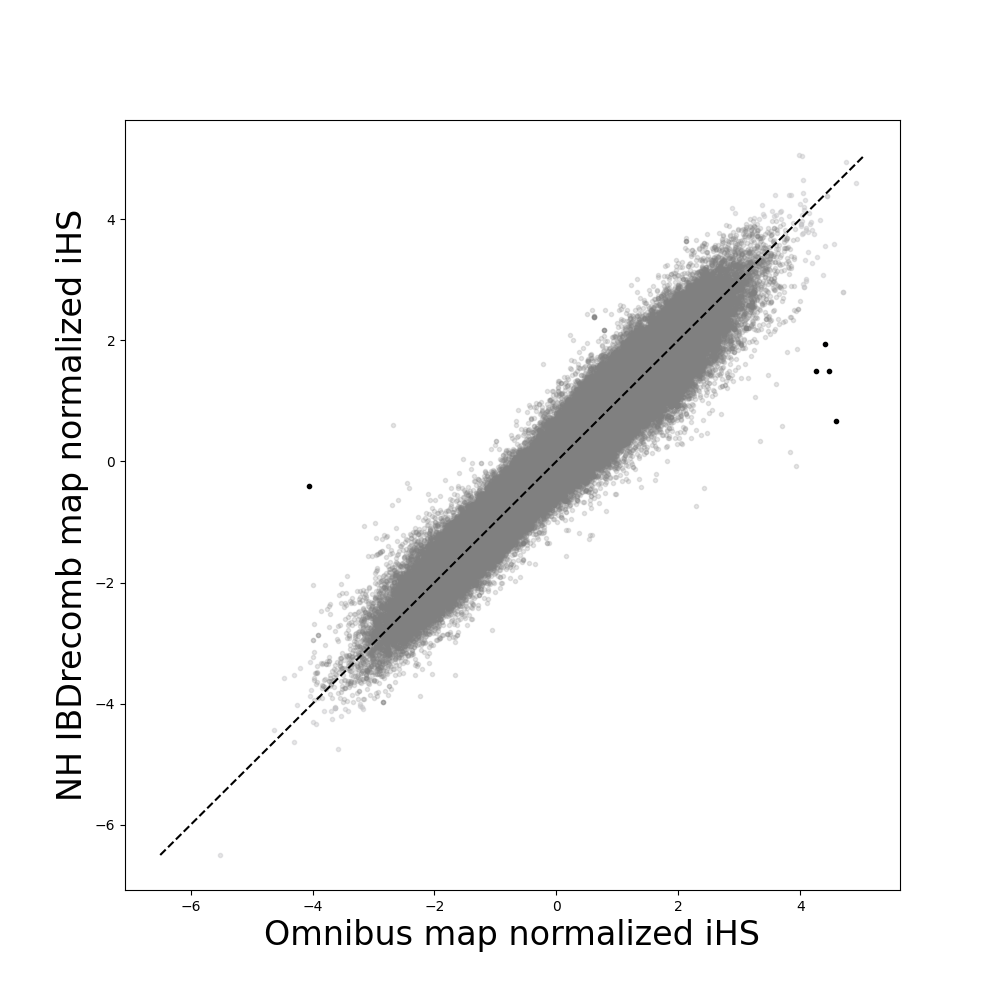
